## Supplementary figures and images for "Rapid progression is associated with lymphoid follicle dysfunction in SIV-infected infant rhesus macaques"

### Supplemental Figure 1

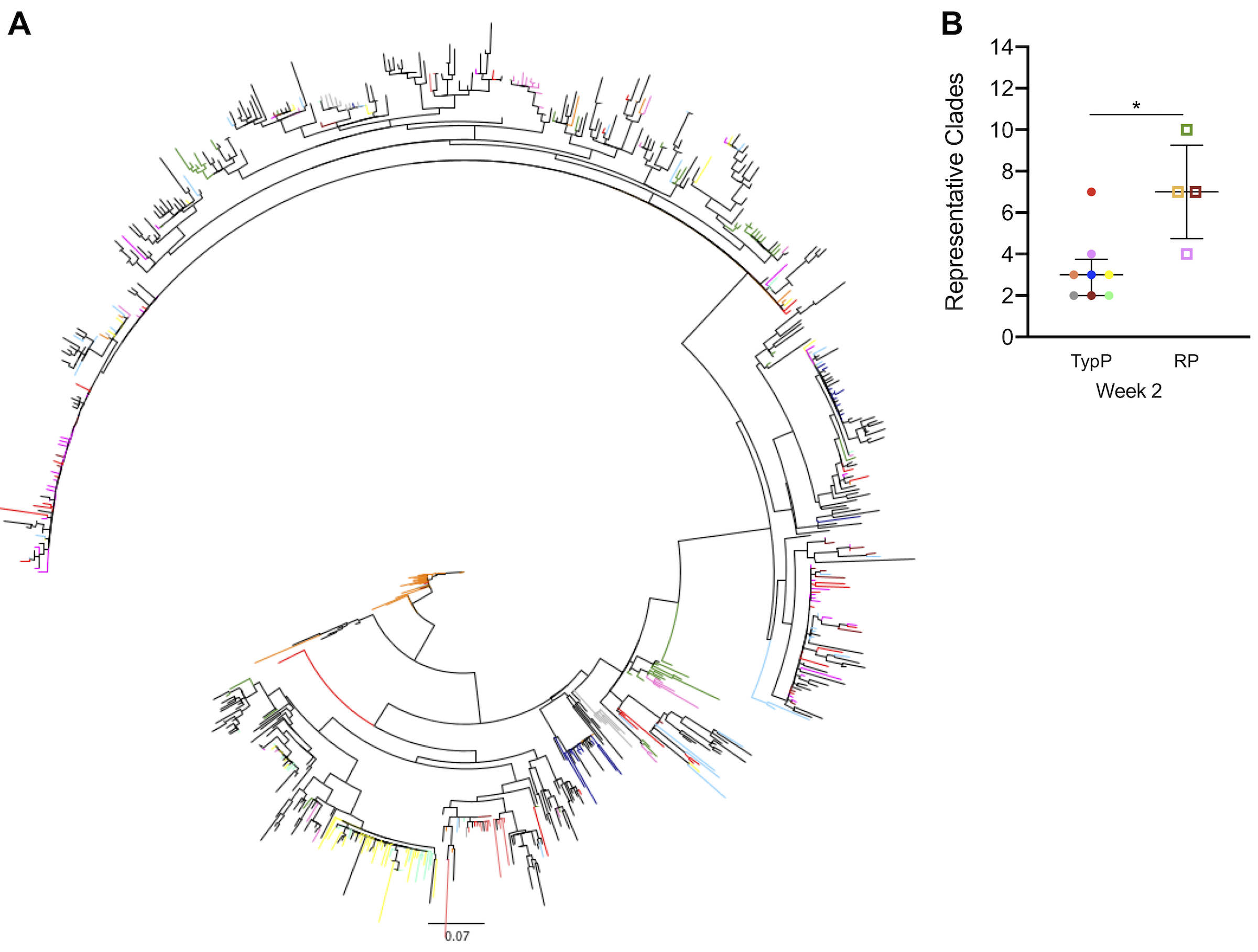

### Supplemental Figure 2

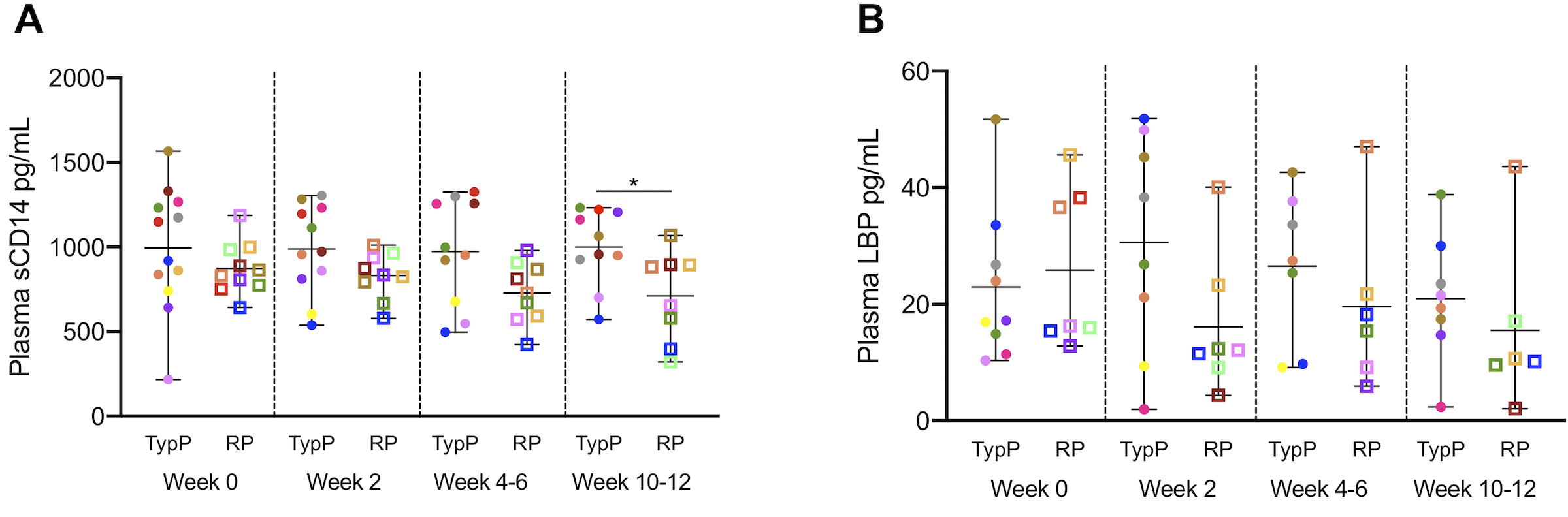

### Supplemental Figure 3

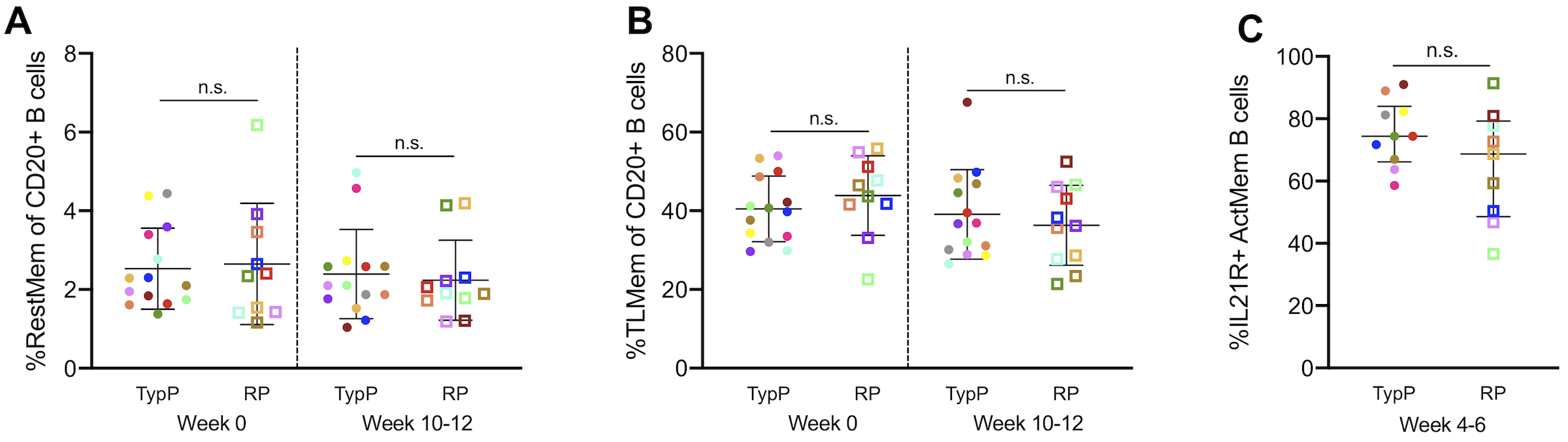
